# Sequence of a *Coxiella* endosymbiont of the tick *Amblyomma nuttalli* suggests a pattern of convergent genome reduction in the *Coxiella* genus

**DOI:** 10.1101/2020.07.17.208157

**Authors:** Tiago Nardi, Emanuela Olivieri, Edward Kariuki, Davide Sassera, Michele Castelli

## Abstract

Ticks require bacterial symbionts for the provision of necessary compounds that are absent in their hematophagous diet. Such symbionts are frequently vertically transmitted and, most commonly, belong to the *Coxiella* genus, which also includes the human pathogen *Coxiella burnetii*. This genus can be divided in four main clades, presenting partial but incomplete co-cladogenesis with the tick hosts. Here we report the genome sequence of a novel *Coxiella*, endosymbiont of the African tick *Amblyomma nuttalli*, and the ensuing comparative analyses. Its size (∼1 Mb) is intermediate between symbionts of *Rhipicephalus* species and other *Amblyomma* species. Phylogenetic analyses show that the novel sequence is the first genome of the B clade, the only one for which no genomes were previously available. Accordingly, it allows to draw an enhanced scenario of the evolution of the genus, one of parallel genome reduction of different endosymbiont lineages, which are now at different stages of reduction from a more versatile ancestor. Gene content comparison allows to infer that the ancestor could be reminiscent of *Coxiella burnetii*. Interestingly, the convergent loss of mismatch repair could have been a major driver of such reductive evolution. Predicted metabolic profiles are rather homogenous among *Coxiella* endosymbionts, in particular vitamin biosynthesis, consistently with a host-supportive role. Concurrently, similarities among *Coxiella* endosymbionts according to host genus and despite phylogenetic unrelatedness hint at possible host-dependent effects.

**Significance statement:** The genus *Coxiella* includes the pathogen *Coxiella burnetii* and widespread nutritional mutualists in ticks. Current knowledge on their evolution is hampered by the limited genomic resources available.

Here we provide the first genome sequence of a *Coxiella* endosymbiont of clade B, the only clade for which none was available.

These data allow to infer an evolutionary scenario of parallel genome reduction among *Coxiella* endosymbionts, with similar constraints, leading to selective retention of biosynthetic pathways beneficial for the host. The combined predicted functional capabilities of the symbionts appear to be a subset of those of *C. burnetii*. Accordingly, this pathogen could be closer to an ancestral state of the endosymbionts, rather than being derived from an endosymbiotic ancestor, as previously hypothesized.

## Introduction

Mutualistic associations with bacteria are widespread and can allow eukaryotes to colonise novel ecological niches (Bennett & Moran 2015). In arthropods, intracellular and maternally transmitted bacterial mutualists are particularly common (Douglas 1998). A typical role of these endosymbionts is providing essential nutrients that are absent in the host’s diet (Sandström & Moran 1999). The classical example is that of sap-feeding insects, such as aphids, acquiring essential amino acids and vitamins from intracellular bacteria (Wernegreen 2012).

Similarly, blood-feeding arthropods also rely on an incomplete source of nutrients and harbour bacterial mutualists. For instance, “*Candidatus* Riesia” bacteria supply their host, the human lice, with several compounds, in particular B vitamins (Boyd et al. 2017; Sasaki-Fukatsu et al. 2006). A better understanding of these symbioses and their role can foster the development of control strategies for hematophagous arthropod vectors of diseases (Zindel et al. 2011). Ticks in particular are extensively studied due to their prominent role of disease vectors for humans and domestic animals (Dantas-Torres et al. 2012). The most important pathogens vectored by ticks include *Borrelia*, the tick-borne encephalitis virus, *Coxiella burnetii*, and multiple *Rickettsiales* bacteria (Kernif et al. 2016).

Most tick species present at least one bacterial endosymbiont, and many species more than one (Cafiso et al. 2016; Moutailler et al. 2016; Duron et al. 2017). Transmission of symbionts is mainly dependent on maternal inheritance through transovarial transmission, but horizontal transfer, possibly during co-feeding, also plays a role. Indeed, closely related tick species, and even different individuals of the same species, may harbour different sets of bacterial symbionts (Duron et al. 2017).

Experimental evidence indicates a major role of such symbionts in tick physiology, as their depletion resulted in impaired growth and reproduction in multiple species belonging to different genera (Zhong et al. 2007; Guizzo et al. 2017; Ben-Yosef et al. 2020). Interestingly, in *Ornithodoros moubata* such developmental defects were rescued by supplementation with B vitamins (Duron et al. 2018). Collectively, these data suggest a common role as B vitamins providers of different, phylogenetically unrelated, symbionts. Other parallel roles have been hypothesized, including protection from pathogens, provision of energy, support for feeding, protection against oxidative and osmotic stress, and waste molecule recycling (Buysse et al. 2019; Olivieri et al. 2019).

Most of characterized tick symbionts are affiliated to *Coxiella* (*Gammaproteobacteria*) (Duron et al. 2017), and will now be abbreviated as CEs (*Coxiella* endosymbionts). Besides many CEs of ticks, this genus includes the tick-borne pathogen *Coxiella burnetii*, causative agent of the Q-fever (Angelakis & Raoult 2010). It has been hypothesized that the latter arose from a CE ancestor, but the origin of its virulence is unclear (Duron et al. 2015).

Currently, a total of seven genomes of CEs are available (Gottlieb et al. 2015; Smith et al. 2015; Guizzo et al. 2017; Ramaiah & Dasch 2018; Tsementzi et al. 2018). These present the hallmarks of genome reduction in bacterial symbionts (McCutcheon & Moran 2012), but at different stages. CEs of *Rhipicephalus* spp. have comparatively larger genomes (1.2-1.7 Mb) with a higher number of pseudogenes (Gottlieb et al. 2015; Guizzo et al. 2017; Ramaiah & Dasch 2018; Tsementzi et al. 2018), characteristics of relatively recent symbioses with ongoing genome reduction. Conversely, the CEs of *Amblyomma* ticks have more streamlined genomes (0.6-0.7 Mb), with high gene density and low number of mobile elements (Smith et al. 2015), typical of a later stage of symbiosis.

Nevertheless, all CEs possess the pathways for production of many B vitamins, consistently with the hypothesised role and the general trend observed in nutritional symbionts, which retain some host-supportive pathways even in case of severe genome reduction (Nakabachi et al. 2006; López-Madrigal et al. 2011).

Four phylogenetic clades (A-D) were identified in the genus *Coxiella*, exhibiting only partial congruence with their hosts’ phylogeny (partial co-cladogenesis). Clade C displays a good degree of co-cladogenesis with *Rhipicephalus* hosts, while CEs of two unrelated clades (B and D) were found in *Amblyomma* hosts (Duron et al. 2015). Interestingly, the known clade B *Amblyomma* hosts came from the African continent, while clade D hosts are American (Duron et al. 2015; Binetruy et al. 2020). Currently, genomes of representatives of three of the four *Coxiella* clades have been sequenced, i.e. CEs from clades C and D, and *Coxiella burnetii* from clade A.

Here, we sequenced the genome of a novel CE of *Amblyomma nuttalli* belonging to the fourth clade (B), and used it for comparative analyses, providing a basis for improving the understanding of the diversity and evolution of CEs.

## Materials and methods

An adult female of *Amblyomma nuttalli* was collected from a white rhinoceros in the Masai Mara National Reserve, Kenya in February 2016. The tick was morphologically identified following standard taxonomic keys (Theiler & Salisbury 1959) and subjected to DNA extraction, using NucleoSpin® Tissue Kit (Macherey Nagel, Duren, Germany), according to the manufacturer’s instructions. DNA was subjected to Illumina HiSeq X by Admera Health (South Plainfield, NJ, USA) using a Nextera XT library, obtaining 27,3511,224 150-nt paired-end reads.

The reads were assembled using SPAdes (3.6.0), and subjected to a modified version of the blobology pipeline (Kumar et al. 2013), in order to select only the symbiont sequences (for a complete description of the process see (Castelli et al. 2019)). Briefly, we selected contigs with a log10 coverage higher than 2.5, extracted and reassembled separately the reads mapping on those contigs (Langmead & Salzberg 2012), and extensively revised manually the results (Supplementary Figure 1-2).

The completeness level of the genome was confronted with all published CE genomes using BUSCO, using the *Gammaproteobacteria* lineage dataset (Seppey et al. 2019).

Genome annotation was performed using Prokka 1.11 (Seemann 2014) and manually curated by inspecting blastp hits of predicted ORFs on NCBI nr, Uniprot, and *Legionellales* sequences.

ISEScan (Xie & Tang 2017) and ISfinder (Siguier et al. 2006) were used to identify insertion sequences and PHASTER (Arndt et al. 2016) for prophages. Pseudogene prediction on the novel genome, all published CE genomes, and representative *C. burnetii* genomes (Supplementary table 1) was performed using Pseudo-finder (Syberg-Olsen & Husnik 2018). COGs were predicted on the same dataset using the NCBI pipeline (Galperin et al. 2015) on validated genes (i.e. ORFs excluding predicted pseudogenes). COG repertoires were used for comparative analyses. Metabolic pathways were manually reconstructed employing the BioCyc database reference (Karp et al. 2019).

Two datasets were used for phylogeny. The first one involved a wide taxonomic sampling, analysed through MLST (multilocus sequence typing) as in (Duron et al. 2015), thus employing five genes and 96 organisms (published dataset plus all available CEs).

The second set was analysed by using a phylogenomic approach, and included the previous selection of *Coxiella* genomes, a representative selection of *Coxiellaceae*, including 1 MAG (metagenome assembled genome), and two other *Legionellales* as outgroup (Supplementary table 1). Using OrthoFinder (2.3.3) (Emms & Kelly 2019), 213 single copy conserved orthologs were identified.

Then, for the two sets, respectively the nucleotide and protein sequences of each single gene were aligned separately using Muscle (Edgar 2004), polished with Gblocks (Talavera & Castresana 2007), and finally concatenated (3,118 and 59,256 total positions, respectively). For each set, we inferred the best model (GTR+I+G and LG+I+G, respectively) using modeltest-ng 0.1.3 (Darriba et al. 2019), built a maximum likelihood tree with RAxML 8.2.4 (Stamatakis 2014) with 1000 bootstrap pseudo-replicates, and a Bayesian inference tree with MrBayes (Ronquist et al. 2012) using three independent runs for 1 million and 250,000 generations, respectively, with a burn-in of 25%.

## Results and discussion

The obtained genome assembly of the CE of *A. nuttalli* has a total length of 1,001,386 bp (9 contigs; N50: 229,733 bp; GC: 35,95%). BUSCO completeness score was 79.2%, similar to other CEs (Supplementary Figure 3). A total of 45 RNA genes (including 38 tRNAs and 3 rRNAs) and 730 ORFs were found. Among these, we identified 696 functional CDSs and 34 pseudogenes, accounting for a total of 658,600 bp (65.7%) coding (including structural RNA genes). Neither prophages nor ISs were found.

MLST phylogeny provided an overall consistent topology with most previous studies (Duron et al. 2015; Gottlieb et al. 2015), in particular for the major *Coxiella* clades and their relationships (clade A earliest divergent, clade B sister group of clades C+D), with moderate to high support (Figure 1A, Supplementary Figure 4-5). Consistently with what could be expected based on geographical origin, the CE of *A. nuttalli* lies in the clade B, composed by CEs of African ticks.

**Fig. 1:**
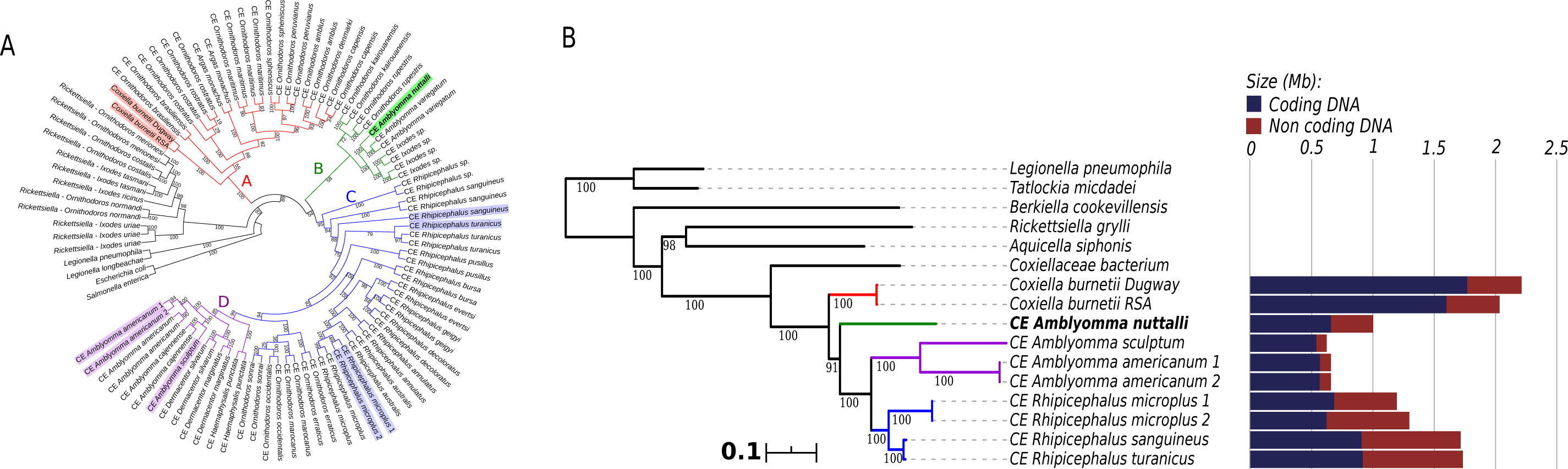
(a) Maximum likelihood MLST phylogenetic tree of *Coxiella* and other *Legionellales* (full tree in Supplementary Figure 4), and (b) maximum likelihood phylogenomic tree of *Coxiellaceae*. In (a,b), the four major *Coxiella* clades are evidenced by different colors, numbers on branches stand for bootstrap support after 1000 pseudo-replicates, CE stands for *Coxiella* endosymbiont, and respective host organisms are indicated. In (a), symbionts with available genomes are highlighted. In (b), scale bar stands for estimated proportional sequence divergence, and the bar plot on the right shows the respective coding (blue) and non-coding (red) genome sizes.

For the available *Coxiella* genomes, phylogenomic analysis showed the same relations of the MLST phylogeny (Figure 1B, Supplementary Figure 6). Interestingly, for all *Coxiella*, including the novel CE of *A. nuttalli*, branch lengths are proportional to the degree of genome reduction (Supplementary Table 2), consistently with previous analyses (Duron et al. 2015; Gottlieb et al. 2015). This would indicate a higher evolutionary rate in smaller genomes, as predicted for obligate symbionts by genome reduction models (McCutcheon & Moran 2012). Specifically, *C. burnetii*, a pathogen capable of living in different environments and hosts (Angelakis & Raoult 2010), presents the largest genome (2 Mb) and, with a high coding density (79.9%), the highest amount of coding DNA (1.8 Mb). All the host-restricted CEs have smaller genomes, with similar sizes within each clade: 1.2-1.7 Mb for CEs of *Rhipicephalus* (clade C), 1.0 Mb for the novel CE of *A. nuttalli* (clade B), and 0.6-0.7 Mb for CEs of other *Amblyomma* species (clade D). However, the degree of genome reduction does not correlate with the phylogenetic branching pattern between clades, in particular the CEs with more reduced genomes (clades B, D) do not form a single monophyletic group (Figure 1B). Accordingly, considering also the novel CE of *A. nuttalli*, a scenario with parallel independent genome reduction in genus *Coxiella* (at least for monophyletic CEs of B-D clades together) appears plausible.

Interestingly, most size variation resides in the non-coding genome (from 88 Kb to 816 Kb), while the length of the functional coding genome is overall less variable in CEs, ranging from 540 Kb of CE of *A. sculptum* (Clade D), to 917 Kb of CE of *R. turanicus* (Clade C). These features as well are consistent with a recent and still ongoing parallel genome reduction of CEs under similar constraints, possibly due to an equivalent role for the host. Accordingly, the CE of *A. nuttalli* has traits of a relatively long-term obligate symbiont, at an intermediate stage among CEs for its genome size and coding density (Supplementary Table 2), and having no predicted mobile elements.

The functional capabilities, as represented by COG repertoires, are consistent with the observed pattern of genome reduction (Figure 2A). *C. burnetii* is the richest in COGs for all functional categories. CEs have lower COGs numbers, roughly proportional to the respective coding genomes. Many core functions are highly conserved, such as translation machinery (J), coenzyme (H) and nucleotide (F) metabolism, energy production (C), protein modification and chaperones (O), lipid synthesis (I), cell cycle regulation (D). This is consistent with their expected major role for bacterial survival and/or host-support (coenzymes). On the other side, all CEs are more pronouncedly depleted in accessory and regulative functions, including poorly characterized ones (R and S), signal transduction (T), secondary metabolite metabolism (Q), cell motility (N), secretion systems (U), extracellular and defense structures (W and V). Such functionalities are probably less important in strictly host-associated bacteria. Notable is the case of type IV secretion, probably ancestral in *Legionellales* (Hugoson et al. 2019) and an important virulence factor in *C. burnetii* (Luedtke et al. 2017), but absent in all CEs. Some functions display gradients of conservation along the genome size, e.g. membrane structure biogenesis (M), which correlates with decrease in lipopolysaccharide complexity, while peptidoglycan synthesis is conserved.

**Figure 2.**
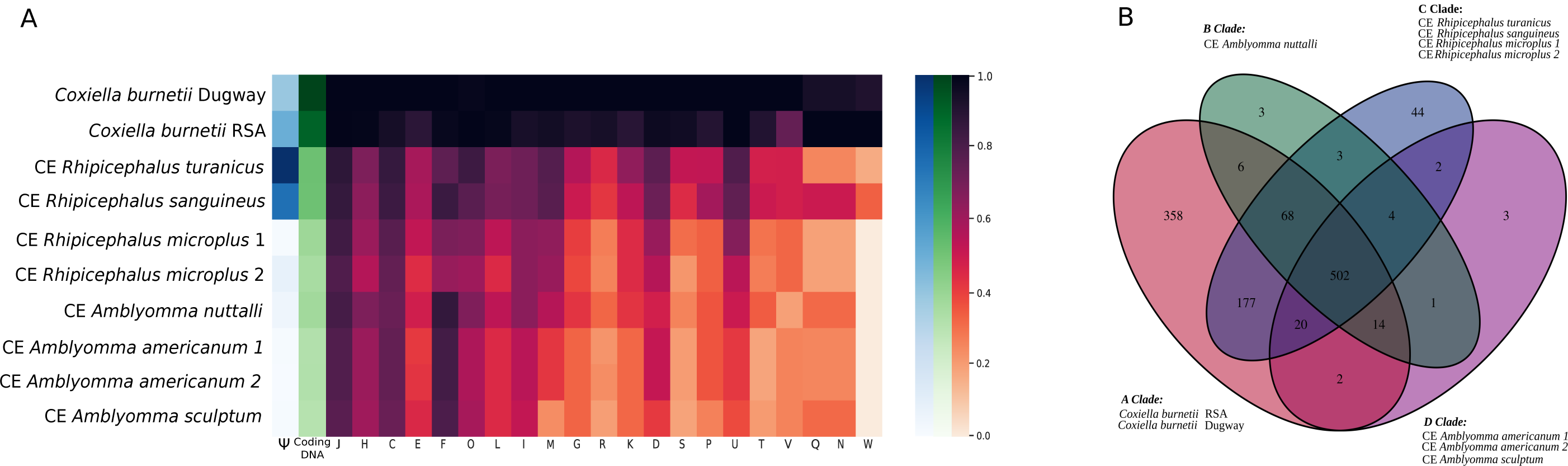
(a) Heatmap showing variations in number of pseudogenes (blue, indicated with the Ψ), genome size (green) and COG (Clusters of Orthologous Groups) repertoire for each), genome size (green) and COG (Clusters of Orthologous Groups) repertoire for each functional category (orange-purple) in the *Coxiella* genus. CE stands for *Coxiella* endosymbiont. Organisms are grouped according to phylogenetic clades, in turn sorted according to coding DNA size. The color intensity is independently scaled for each column in proportion to its maximum value. (b) Venn diagram representing COG distribution on *Coxiella* clades. For each clade, COGs identified in at least one of the listed genomes are counted.

Such scenario is reflected at the level of single COGs (Figure 2B), as *C. burnetii* (clade A) presents the highest number of unique COGs. CEs of all clades are a substantial subset of *C. burnetii*, which lacks only 60 of the 1207 total COGs in the dataset. Similar observations can be drawn among progressively more reduced CE clades, with highly streamlined clade D CEs almost as a subset of the other CEs.

Few lineage-specific peculiarities were found, and the functional significance of many of these is unclear, e.g. the *A. nuttalli*-specific COGs display redundant or poorly characterized functions (Supplementary table 3). However, some relevant variability was also observed. All CEs retain the predicted capability to perform glycolysis, Krebs cycle and oxidative phosphorylation (Supplementary table 4). However, contrarily to the other CEs, the symbiont of *A. nuttalli* does not possess a conventional citrate synthase, but instead presents a 2-methylcitrate synthase, which may also catalyse the same reaction (Patton et al. 1993). As *C. burnetii* has both genes, this partial redundancy could have been ancestral, and independently lost in different CE lineages.

In general, while biosynthetic abilities for amino acids are scarce in all *Coxiella* (E), those for vitamins and cofactors (H) are, as expected, abundant and highly conserved (Supplementary table 4). CEs are in particular rich in genes for the synthesis for riboflavin (B2), pantothenate (B5) and its derivative CoA, pyridoxine (B6), folic acid (B9), and biotin. For biotin (and lipid) synthesis, missing FabI functionality is possibly replaced by FabV (Massengo-Tiassé & Cronan 2008). Interestingly, despite overall smaller functional capabilities, the CEs of *Amblyomma* display complete biosynthetic pathways for thiamine (B1) and NAD (B3) (except the last thiamine step in clade D), while symbionts of *Rhipicephalus* retain only partial pathways (including final steps from thiamine phosphate and β-nicotinate D--nicotinate D-ribonucleotide). Such differences can be explained by the presence of not yet identified transporters and/or non-canonical enzymes (Gottlieb et al. 2015). They might also indicate different, not yet clarified, metabolic requirements of the tick hosts, with *Amblyomma* species requiring a full pathway while *Rhipicephalus* being permissive for the loss of some genes.

A similar scenario may hold for nucleotide metabolism (F), which is also more reduced in the CEs of *Rhipicephalus*, lacking the initial path for the synthesis of purines (up to 5-aminoimidazole ribonucleotide) (Supplementary table 4).

Consistently with their symbiotic condition (McCutcheon & Moran 2012), CEs are depleted in DNA repair abilities (L), with lineage-specific features (Supplementary table 4). For example both CEs of *R. microplus* are devoid of the RecFOR pathway and RecA, involved in homologous recombination (Kuzminov 1999). Interestingly, the MutSL pathway is fully absent in the smaller genomes of CEs of *Amblyomma* of clades B and D, but complete in *C. burnetii* (clade A) and in most members of clade C. Among those, the exception is the CE of *R. microplus* “2”, found to have a full *mutL* gene, but a truncated *mutS* pseudogene, probably retaining only partial or no functionality. Parsimoniously, we can identify at least three multiple convergent losses of this pathway among CEs (complete in the clades B and D, and still ongoing within clade C). Considering the strong correlation with the degree of genome reduction, it is reasonable to hypothesise that this loss may have had a major evolutionary impact, possibly directly causing increased mutation rates (Schofield & Hsieh 2003), and eventually resulting in accelerated and more pronounced genome reduction. Thus, the seminal speculations by Gottlieb and co-authors (2015) on a smaller dataset are reinforced. This effect would be particularly evident from the differences in coding size and functional categories among the two closely related CEs of *R. microplus* (Figure 1B).

## Conclusions

The novel sequence of CE of *Amblyomma nuttalli* expands the available diversity of *Coxiella* genomes, being the first obtained from clade B. Despite reduced genome size, biosynthetic pathways for vitamins appear to be conserved, as in other CEs, supporting a role of CEs in dietary supplementation of these compounds to the hosts. At same time, some variations were found in vitamin and purine synthesis, possibly dependent on the host species.

Combining phylogenetic and genomic data, an evolutionary scenario of parallel genome reduction with analogous constraints among CEs (clades B-D) can be drawn. Consistently with previous observations (Gottlieb et al. 2015), the convergent loss of MutSL could have been a driver of such reduction, with the CE of *A. nuttalli* representing an intermediate level between clade C and D, and all CEs substantially being a subset of *C. burnetii*. Accordingly, and differently from previous views (Duron et al. 2015), CEs could have evolved from a more versatile *C. burnetii*-like ancestor, analogously to other unrelated symbionts (Taylor et al. 2005; Gerhart et al. 2016). Genomic and phylogenomic analyses of representatives of clade A other than *C. burnetii* may provide further insights.

## Data Availability Statements

The data underlying this article are available in the NCBI GenBank Database at ncbi.nlm.nih.gov/, and can be accessed with JACBPR000000000 (CE genome sequence) and with SRR12168527 (total reads in SRA).

## Acknowledgements

This work was supported by the Human Frontier Science Programme Grant RGY0075/2017 to DS, and by the Italian Ministry of Education, University and Research (MIUR): Dipartimenti di Eccellenza Programme (2018–2022)—Department of Biology and Biotechnology “L. Spallanzani”, University of Pavia to DS.

