## Supplementary material for "Sequence of a *Coxiella* endosymbiont of the tick *Amblyomma nuttalli* suggests a pattern of convergent genome reduction in the *Coxiella* genus": Legend of supplementary materials

**Supplementary Table legends**

Supplementary Table S1: Genomes employed in this study. For phylogenomic all were used, for functional and comparative analyses only those highlighted in bold (*Coxiella* genus).

Supplementary Table S2: Total and coding genome size in *Coxiella* bacteria.

Supplementary Table S3: COGs of the CE of *Amblyomma nuttalli*, divided into the peculiar ones and those shared with other *Coxiella*.

Supplementary Table S4: Gene presence/absence profile for selected pathway within genus *Coxiella*.

**Supplementary Figure legends**

Supplementary Figure S1: Blobology plot of the preliminary assembly: each dot represents a contig, colored according to the taxonomy of its best megablast hit. The x and y axes represent, respectively, GC content and log10 of the coverage of the contigs. For readers' clarity, only contigs >1000 bp are shown.

Supplementary Figure S2: Bandage graphical representation (Wick et al. 2015) of the 9 contigs of the final assembly of the *Coxiella* endosymbiont of *Amblyomma nuttalli* and their predicted connections in the assembly graph. The respective contig lengths are indicated.

Supplementary Figure S3: Busco output showing the estimated completeness of the *Coxiella* genome assemblies, according to presence of 366 conserved orthologs in *Gammaproteobacteria*.

Supplementary Figure S4: Full representation of Fig. 1A tree (maximum likelihood MSLT phylogenetic tree of *Legionellales*), with branch lengths proportional to estimated sequence divergence. Clades are labeled as in Fig. 1.

Supplementary Figure S5: Bayesian inference MLST phylogenetic tree, using the same dataset of Fig. 1A and S4. Numbers on branches stand for posterior probabilities, and branch lengths are proportional to the estimated sequence divergence. Clades are labeled as in Fig. 1.

Supplementary Figure S6: Bayesian inference phylogenomic tree with the same dataset of Fig. 1B. Numbers on branches stand for posterior probabilities, and branch lengths are proportional to the estimated sequence divergence. Clades are labeled as in Fig. 1.
