## Supplementary figures and images for "Sequence of a *Coxiella* endosymbiont of the tick *Amblyomma nuttalli* suggests a pattern of convergent genome reduction in the *Coxiella* genus"

### Supplementary figure 1

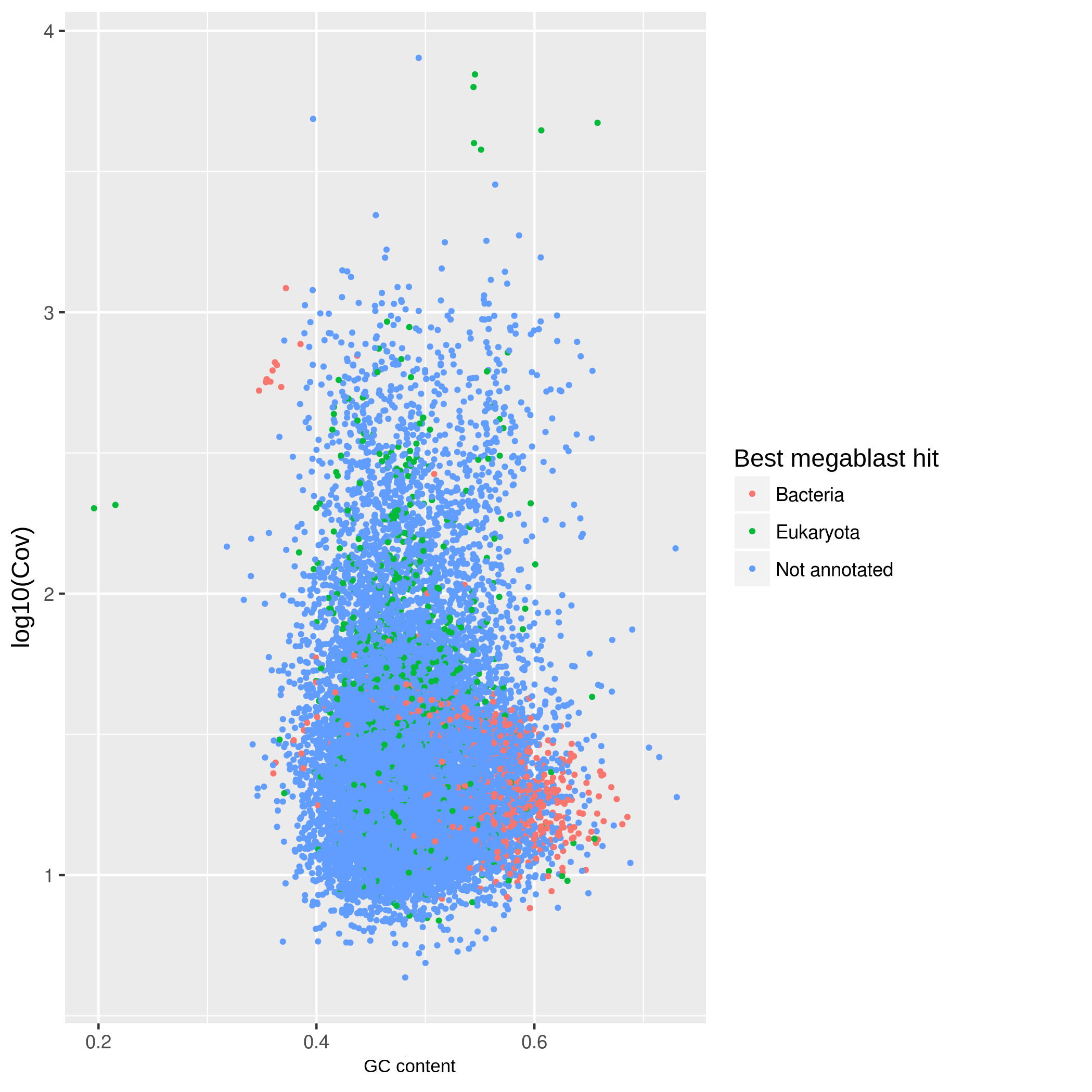

### Supplementary figure 2

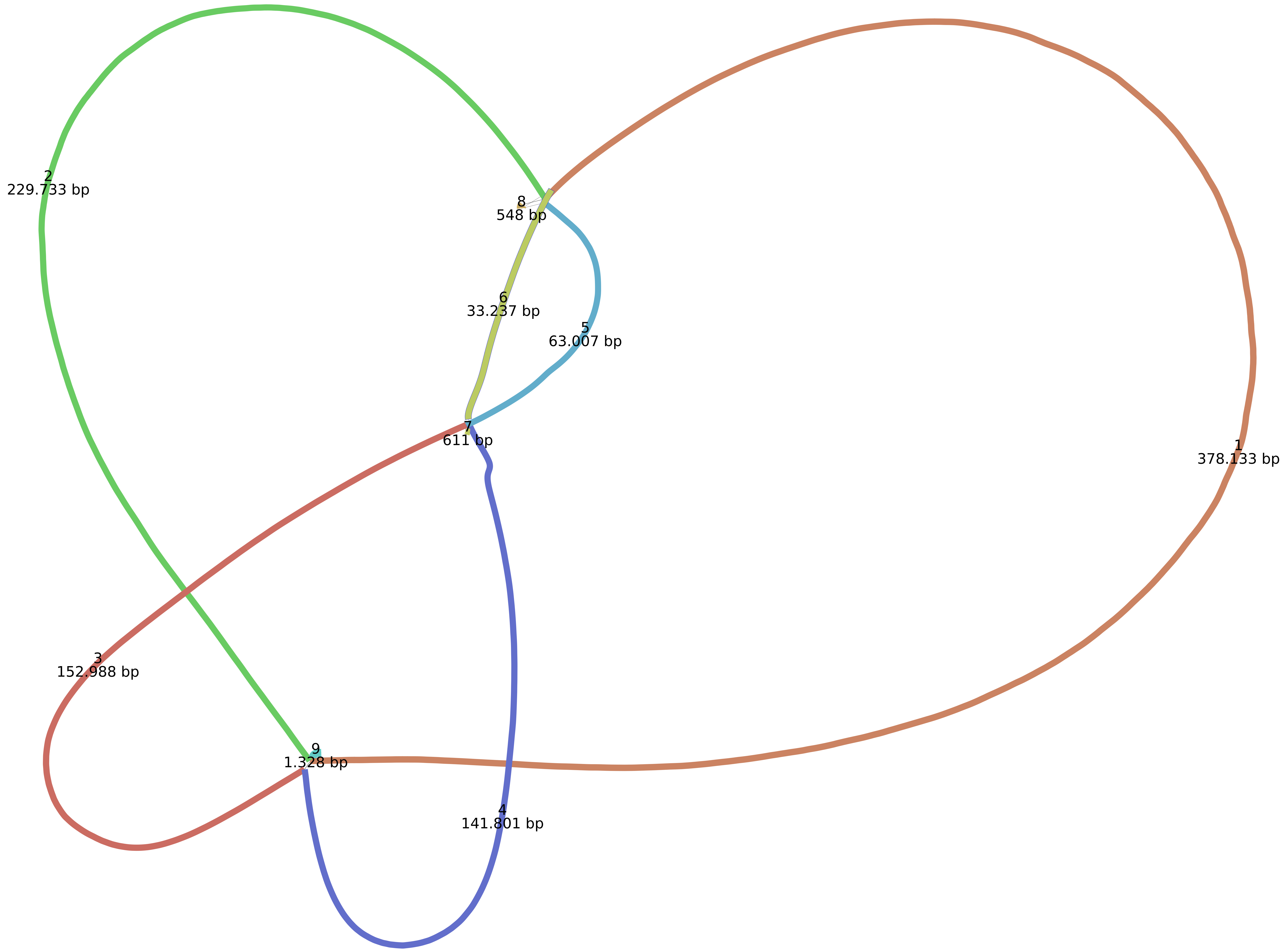

### Supplementary figure 3

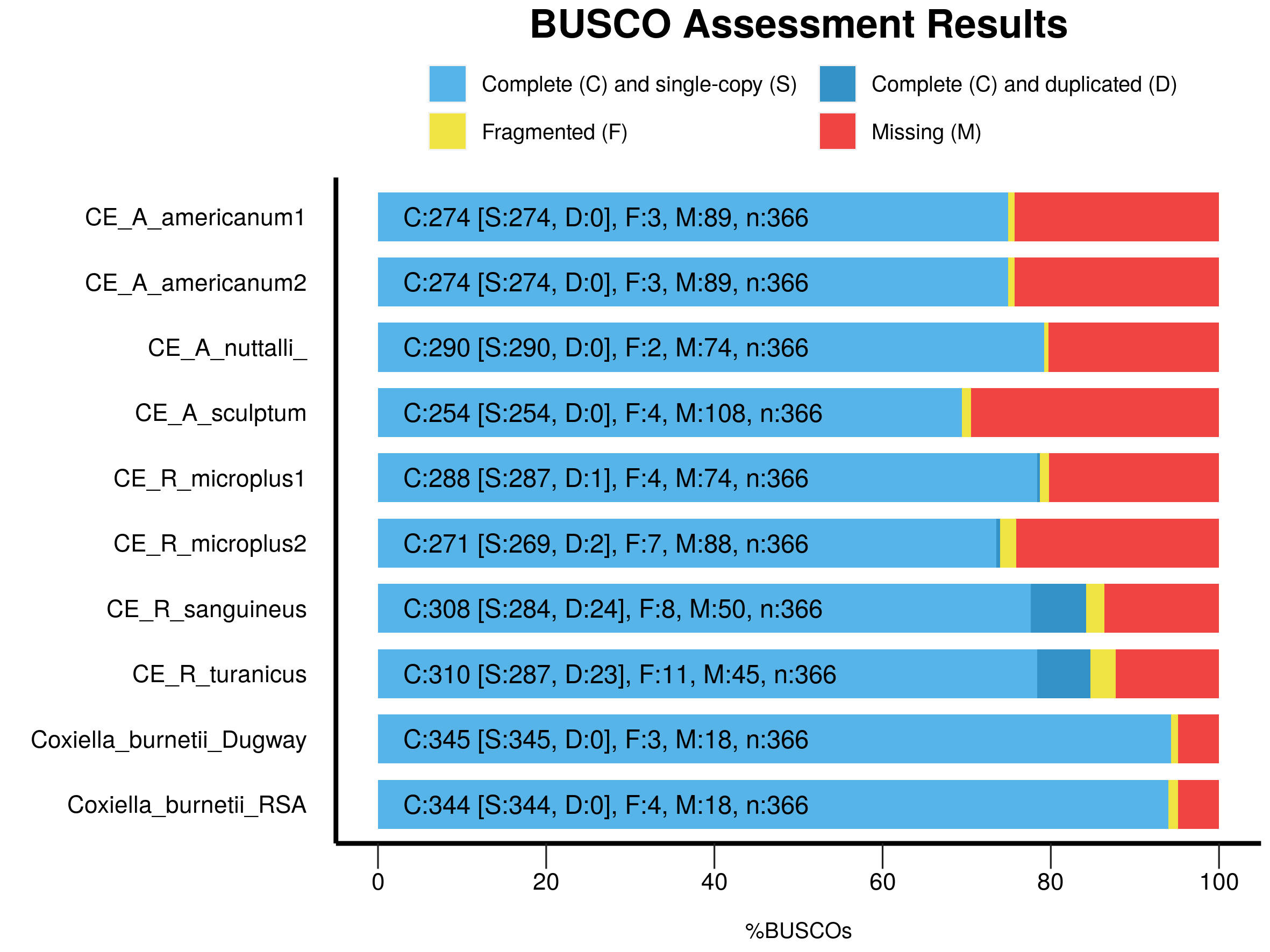

### Supplementary figure 4

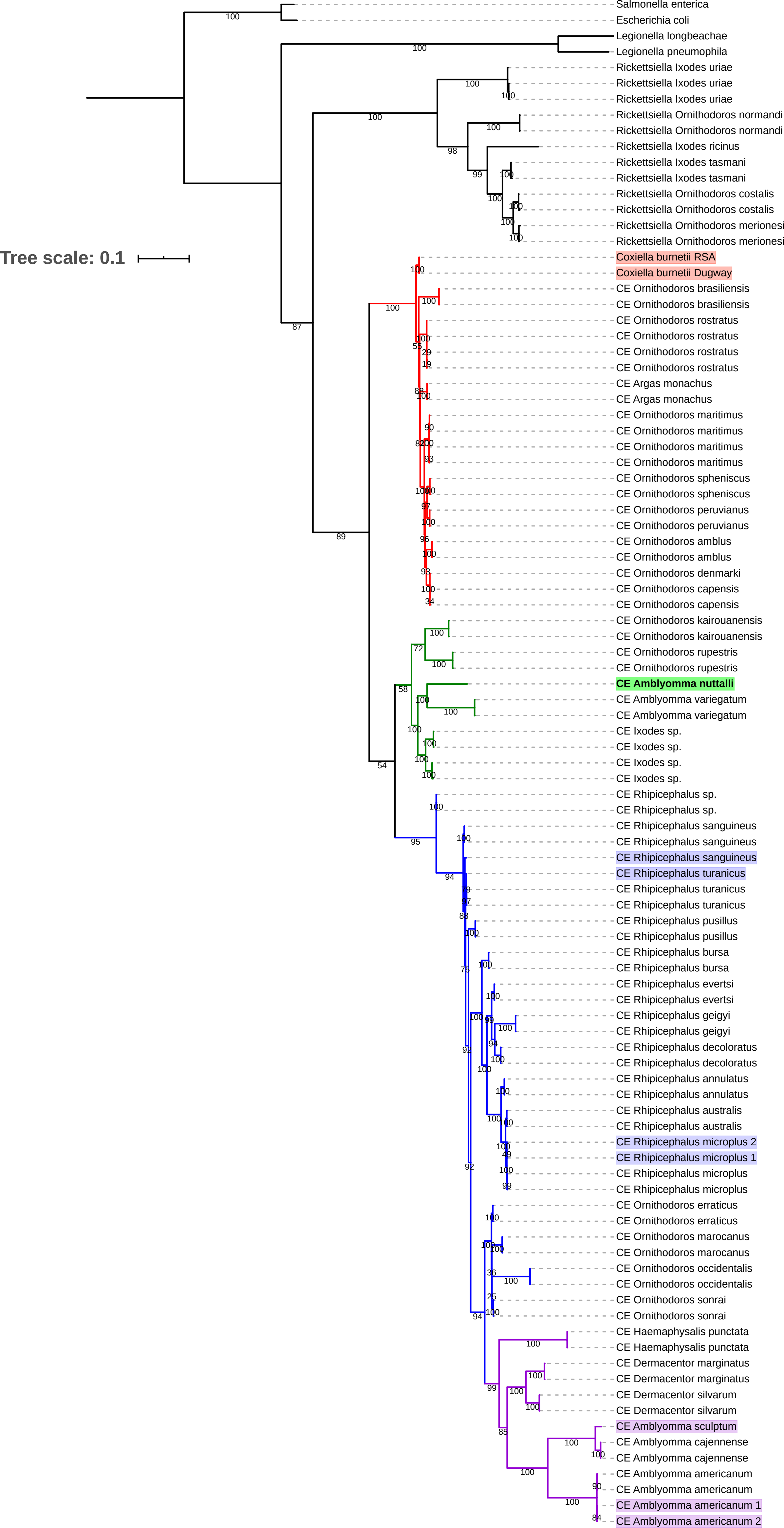

### Supplementary figure 5

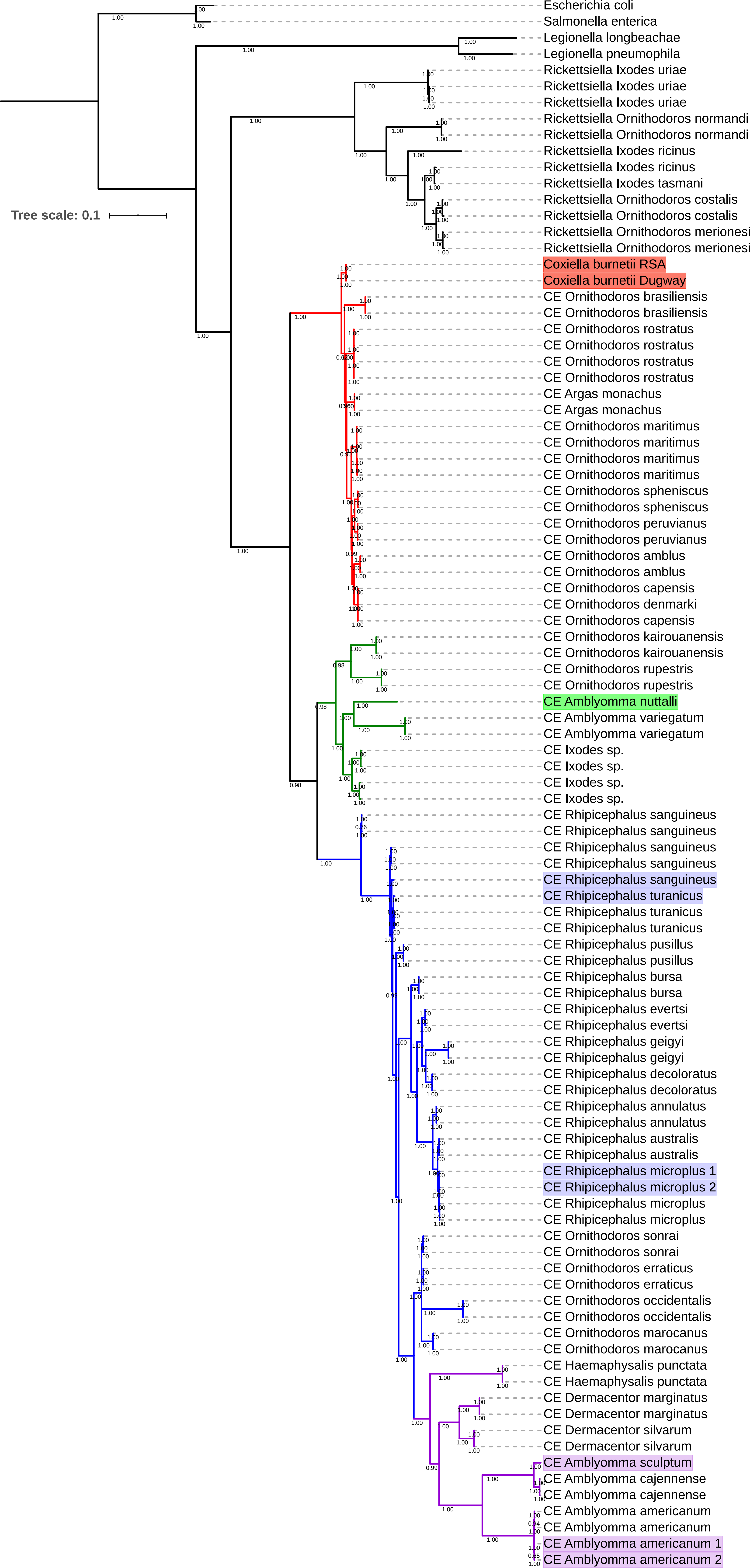

### Supplementary figure 6

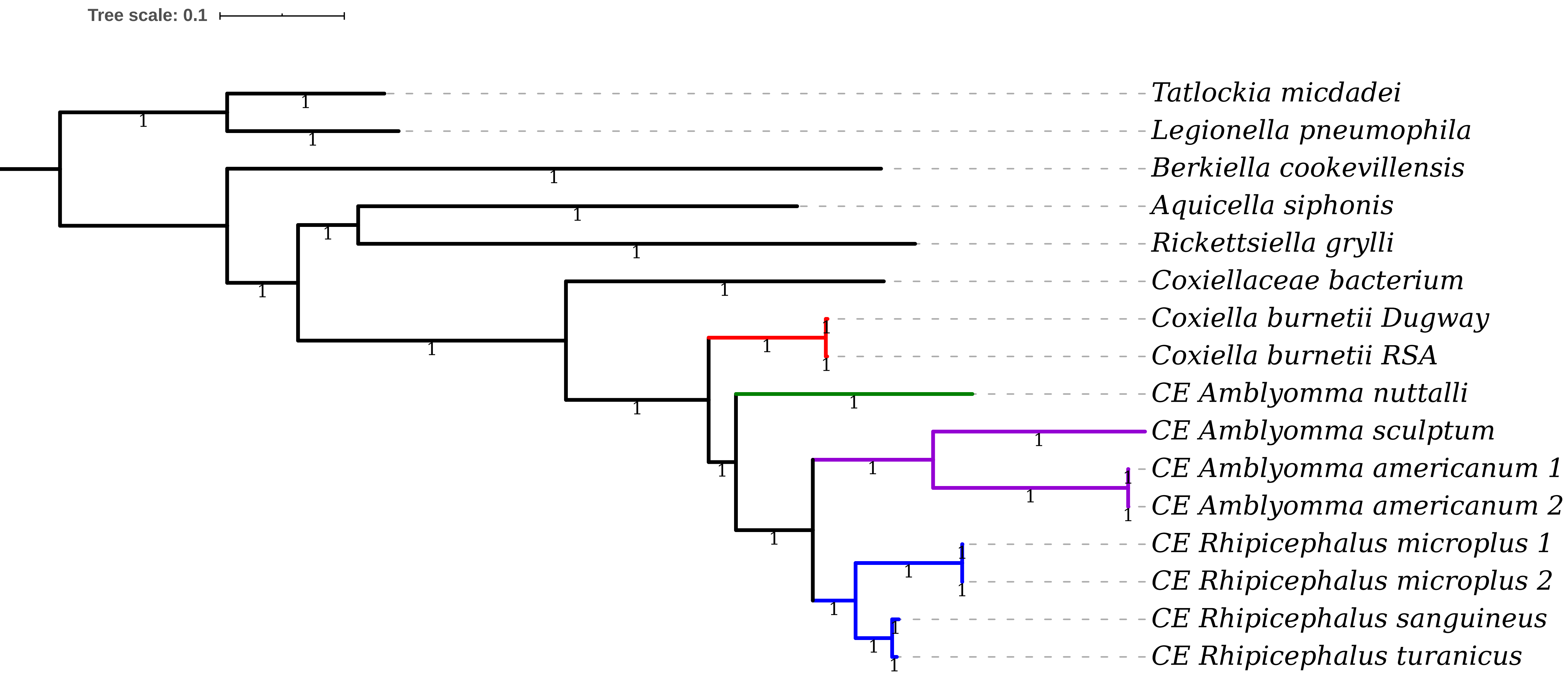
